# Neural encoding of biomechanically (im)possible human movements in occipitotemporal cortex

**DOI:** 10.1101/2025.01.07.631720

**Authors:** Giuseppe Marrazzo, Federico De Martino, Albert Mukovskiy, Martin A. Giese, Beatrice de Gelder

## Abstract

Understanding how the human brain processes body movements is essential for clarifying the mechanisms underlying social cognition and interaction. This study investigates the encoding of biomechanically possible and impossible body movements in occipitotemporal cortex using ultra-high field 7Tesla fMRI. By predicting the response of single voxels to impossible/possible movements using a computational modelling approach, our findings demonstrate that a combination of postural, biomechanical, and categorical features significantly predicts neural responses in the ventral visual cortex, particularly within the extrastriate body area (EBA), underscoring the brain’s sensitivity to biomechanical plausibility. Lastly, these findings highlight the functional heterogeneity of EBA, with specific regions (middle/superior occipital gyri) focusing on detailed biomechanical features and anterior regions (lateral occipital sulcus and inferior temporal gyrus) integrating more abstract, categorical information.

## Introduction

Human bodies convey essential information about others’ actions, intentions, and emotions and provide critical cues in social communication (de Gelder, 2006; de Gelder et al., 2010; Peelen & Downing, 2007; Tipper, Signorini, & Grafton, 2015). Previous research using functional magnetic resonance imaging to investigate the neural basis of body perception (fMRI) has primarily focused on localizing high-level visual category-specific representations. Specific regions in the occipitotemporal and fusiform cortex are selectively responsive to images of bodies, the extrastriate body area (EBA) and the fusiform body area (FBA) (Downing, Jiang, Shuman, & Kanwisher, 2001; Peelen & Downing, 2005). Similar findings of distinct body sensitive patches were found in monkeys in the ventral bank of the superior temporal sulcus (STS), namely the middle STS body patch (MSB) and the anterior STS body patch (ASB), with a putative homology between MSB and EBA, and ASB and FBA (Vogels, 2022). When dynamic images or functional aspects of body perception like action and emotional expression are also considered, body sensitivity was reported in other areas (de Gelder & Poyo Solanas, 2021). This has raised interest in investigating the neural mechanisms underlying body sensitivity, notably in the specific computational mechanisms operating across these different body sensitive areas.

Some studies suggested that EBA is more involved in processing body parts and local features and FBA devoted to holistic processing (Taylor & Downing, 2011; Taylor, Wiggett, & Downing, 2007). There is also some evidence that EBA and FBA might process a combination of local and global body features (Bracci, Ietswaart, Peelen, & Cavina-Pratesi, 2010; Downing & Peelen, 2011, 2016; Marrazzo, De Martino, Lage-Castellanos, Vaessen, & de Gelder, 2023), depending on semantic attributes such as emotion and action (de Gelder, Snyder, Greve, Gerard, & Hadjikhani, 2004; Downing, Peelen, Wiggett, & Tew, 2006; Hadjikhani & de Gelder, 2003), and that EBA is sensitive to task demands (Marrazzo, Vaessen, & de Gelder, 2021). Additionally, recent findings further suggest that activity in the Default Mode Network (DMN) is sensitive to the contrast between biological and non-biological motion based on the naturalness of kinematic patterns. Specifically, the DMN’s stronger response to human-like motion, particularly when it matches expected kinematics, suggests that it may modulate or support EBA and FBA processing by enhancing sensitivity to motion patterns that carry social and biological relevance (E. Dayan et al., 2016).

However, despite these insights, there is no clear understanding of a functional division of labour between different body-sensitive areas. A better understanding of the computational processes within these body-selective areas should clarify their specific contributions to body perception.

Over the past decade, (linearized) encoding (Kay, Naselaris, Prenger, & Gallant, 2008; Naselaris, Kay, Nishimoto, & Gallant, 2011) has been used to compare different computational hypotheses of brain function. In these approaches, brain activity (e.g., blood oxygen level-dependent (BOLD) signals in a voxel or brain region during fMRI) is predicted based on stimulus features derived from computational models. The accuracy of these predictions can then be compared to adjudicate between competing models, or to determine the relative contribution (the variance explained) of each model (Dumoulin & Wandell, 2008; Dupré la Tour, Eickenberg, & Gallant, 2022; Moerel, De Martino, & Formisano, 2012; Nunez-Elizalde, Huth, & Gallant, 2019; Santoro et al., 2014; Thirion et al., 2006; Wandell, Dumoulin, & Brewer, 2007). Encoding models predict neural responses based on specific stimulus features and have been successfully applied to visual processing in early visual cortex (Kay et al., 2008; Naselaris et al., 2011) as well as higher visual cortex (Huth, Nishimoto, Vu, & Gallant, 2012; Marrazzo et al., 2023; Nunez-Elizalde et al., 2019; Yamins et al., 2014). An earlier study used encoding models to human body-selective regions (Marrazzo et al., 2023) and shed light on the relevance of joint positions and their spatial configuration for the responses in the EBA to still images. Like most prior research in the field, the use of still images, only addressed postural aspects rather than movement, thus limiting our understanding of how the brain processes more complex, dynamic information.

Here, we probed EBA’s dependency on joints configuration by using biomechanical manipulations of natural movements based on 3D motion capture (mocap) data. Creating videos that disrupt the natural spatial configuration of joints allowed us to investigate how EBA processes biomechanical plausibility. This approach is particularly important with moving bodies, as dynamic stimuli capture the temporal and kinematic properties essential for understanding how the brain encodes real-world, biologically relevant movements. We specifically tested the hypothesis that EBA is sensitive to biomechanical characteristics of body movements, building on some earlier indications in the literature. For instance, participants exhibit automatic imitation effects even for impossible movements, indicating the brain’s predisposition to process action dynamics despite biomechanical violations (Longo, Kosobud, & Bertenthal, 2008). Recognition of human bodies is significantly affected by inversion, reflecting specialized perceptual mechanisms for recognizing human shape in upright configurations (Reed, Stone, Bozova, & Tanaka, 2003). More recent studies have shown that prior knowledge of biomechanical constraints biases visual memory, with participants misremembering extreme postures as less extreme, adjusting their perceptions toward more biomechanically plausible positions (Han, Gandolfo, & Peelen, 2024). Developmental evidence also points to an early sensitivity to biomechanical constraints on human movement. 12-month-old infants as well as and adults spend more time looking at the elbows during impossible arm movements compared to possible ones (Morita et al., 2012), and newborns can differentiate between biomechanically possible and impossible hand movements (Longhi et al., 2015). Investigating the neural correlates of humanly impossible movements has further revealed that impossible finger movements elicit distinct neural responses compared to possible ones in EBA (Costantini et al., 2005). The influence of biomechanics to the processing of visual information related to the body may be fundamental to how body representations are formed in the brain, and may involve areas like the EBA.

To investigate the computations underlying the neural responses to body movements in the occipitotemporal cortex, we utilized ultra-high-field 7 Tesla fMRI and linearized encoding models, assessing macroscopic and mesoscopic (layer-specific) responses related to biomechanical sensitivity. We aimed to identify how different cortical layers within the EBA encode biomechanical information and distinguish between possible and impossible movements. We employed three distinct encoding models to probe these computations: the 3D Keypoints (kp3d) model, which represents three-dimensional coordinates of body joints and captures precise postural information; the Similarity Distances (simdist) model, which quantifies biomechanical differences between possible (natural) and morphed (impossible) movements based on motion capture data (Ghorbani et al., 2021); and the categorical differences model, which provides a higher-level distinction by categorizing movements as biomechanically possible or impossible. By comparing model performance across cortical layers, we aimed to test the hypothesis that superficial cortical layers encode categorical information—indicating sensitivity to global, higher-order features—while deeper layers encode joint-specific and biomechanical information (contained in the kp3d and simdist models).

## Material and methods

### Participants

12 right-handed subjects (5 males, mean age = 27.8 ± 3.8 years) participated in this study. They all had normal (or corrected to normal) vision and reported no history of psychiatric or neurological disorders. One participant was excluded from the main analysis for excessive head motion across multiple runs. All subjects were naïve to the task and the stimuli and received monetary compensation for their participation. Scanning sessions took place at the neuroimaging facility Scannexus at Maastricht University (NL). All experimental procedure conformed to the Declaration of Helsinki and the study was approved by the Ethics Committee of the faculty of Psychology and Neuroscience of Maastricht University.

### Main experiment stimuli

The stimulus set consisted of 120 videos of two avatars (1 male). The videos were generated by animating mocap data from the MoVi dataset (Ghorbani et al., 2021), which includes recordings from 60 female and 30 male actors performing 21 daily actions and sports movements. For this experiment, we animated six specific actions (kicking, pointing, waving, jumping, jumping jacks, and walking sideways) performed by 17 actors (9 males). The movements of these 17 actors were then used to animate the two avatars, ensuring that the presented stimuli maintained diversity in motion while being standardized in appearance. This process resulted in 96 videos depicting natural body movements. Additionally, we modified the joint angles of the limbs to create 96 biomechanically impossible videos. To refine the set for the fMRI experiment, we conducted a behavioral validation, to select stimuli showing the greatest difference between possible and impossible movements. This ultimately reduced the set to 120 videos (60 possible videos created from 17 actors performing 4 actions: kicking, jumping, pointing, waving). More details are provided in the behavioral validation section below. Each video was edited to have a length between 60 and 90 frames, corresponding to 2 to 3 seconds at 30 frames per second. Additionally, the avatars in each video were aligned to be centered relative to the fixation cross, ensuring a consistent starting position across all videos. During the experiment, the stimuli spanned a mean width and height of 1.84° x 4.32° of visual angle (Fig. 1a).

**Figure 1.**
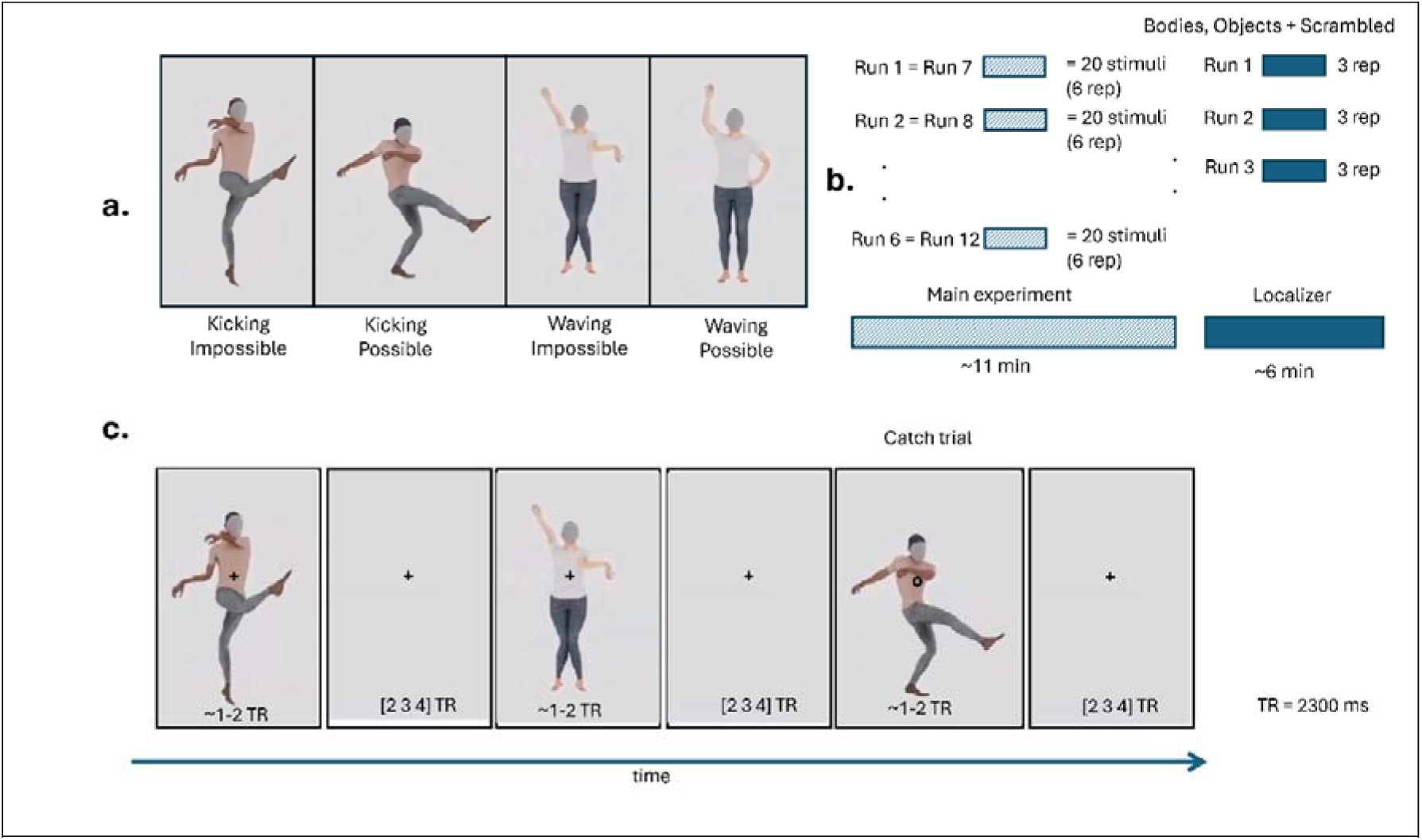
Stimuli and experimental procedure. **(a)** The videos were generated by animating mocap data from the MoVi dataset (Ghorbani et al., 2021). Sixty possible videos were created from 17 actors performing 4 actions: kicking, jumping, pointing, waving. Additionally, we modified the joint angles of the elbows and knees to create 60 biomechanically impossible videos. In panel (a) we show frame of possible videos and their equivalent impossible. **(b)** For each run 1/6 of the stimuli (20) where presented in a pseudo-randomized order following a fast event-related design. Each stimulus was repeated three times per run. Each run was repeated two times across sessions resulting in a total of 120 stimuli repeated six times. To identify body sensitive region, the localizer stimuli included videos of humans performing natural body movement, objects, and their scrambled version. We presented stimuli following a block-design with each block repeated three times per run. **(c)** During the main experiment participants fixated on the cross and were presented with the stimuli depicting possible and impossible body movement for 1-2 TRs (depending on the length of each video) followed by a blank screen which appeared for 2, 3 or 4 TRs. When the fixation cross turned to a circle, they had to press a button whether with the right index finger. TR= 2300ms.

### Localizer stimuli

Stimuli for the localizer experiment consisted of videos depicting two object categories: bodies, objects. Additionally, also a scrambled version of each stimulus was included. (Fig. 1b). The size of the stimuli was 3.5 * 7.5 degrees for human bodies and objects. For more details about the localizer stimuli we refer to (Li et al., 2023). None of the stimuli from the localizer were used in the main experiment.

### Behavioural validation

The stimuli created from the mocap data comprised 96 videos of natural body movements (possible) and their corresponding modified versions, for a total of 192 stimuli. These modified versions (impossible) were created by altering the joint angles of the limbs to produce biomechanically impossible movements. We violated the anatomical constraints of the elbows and knees, by mirroring those joints orientations for each time point of a trajectory. Accordingly, we modified the shoulders and wrist joint angles, as well as ankles and hips, in order to preserve the end-effectors (hands and feet) orientations to be as close as possible to the original (possible) ones for every time point.

Out of the total 192 videos, we selected 120 (60 possible and their impossible version) for the fMRI experiment through a process of behavioral validation. This selection was based on identifying the stimuli that best demonstrated the intended differences between possible and impossible movements, ensuring the most effective set for the experiment. We asked 136 participants (25 males, mean age = 21.45 ± 2 years) to rate the stimuli using a questionnaire consisting of two Likert-scale questions and one categorical question. Participants were presented with half (96) of the total stimuli (192) once. For each participant, the stimuli were pseudo-randomized (96 stimuli randomly selected for each participant, but evenly distributed so that each stimulus was rated by approximately the same number of participants: mean number of responses = 68 ± 2.24). After each presentation, participants were asked to answer a total of three questions about the plausibility/realism of the body movement, action content and salience of specific body parts (see Supplementary materials).

### MRI acquisition and experimental procedure

Participants viewed the stimuli while lying supine in the scanner. Stimuli were presented on a screen positioned behind participant’s head at the end of the scanner bore (distance screen/eye = 99 cm) which the participants could see via a mirror attached to the head coil. The screen had a resolution of 1920x1200 pixels, and its angular size was 16° (horizontal) x 10° (vertical). The experiment was coded in Matlab (v2021b The MathWorks Inc., Natick, MA, USA) using the Psychophysics Toolbox extensions (Brainard, 1997; Kleiner, Brainard, & Pelli, 2007; Pelli, 1997).

Each participant underwent two MRI sessions, we collected a total of twelve functional runs (six runs per session) and one set of anatomical images. Images were acquired in a 7T MR scanner (Siemens Magnetom) using a 32-channel (NOVA) head coil. Anatomical (T1-weighted) images were collected using MP2RAGE MP2RAGE: 0.7 mm isotropic, repetition time (TR) = 5000 ms, echo time (TE) = 2.47 ms, matrix size= 320 x 320, number of slices = 240. The functional dataset (T2*-weighted) covered the occipitotemporal cortex and was acquired using a Multi-Band accelerated 2D-EPI BOLD sequence, multiband acceleration factor = 2, voxel size = 0.8 mm isotropic, TR = 2300 ms, TE = 27 ms, number of slices = 58 without gaps; matrix size = 224 x 224; number of volumes = 300, GRAPPA factor =3. In addition to functional images, phase images were simultaneously acquired along with five noise volumes appended at the end of each run.

During the main experiment, stimuli were presented on the screen for 2-3 seconds (depending on the length of each video) with an inter stimulus interval that was pseudo-randomised to be 2, 3 or 4 TRs. Participants were asked to fixate at all times on a white cross at the centre of the screen (Fig. 1c).

To control for attention, participants were asked to detect a shape change at the fixation cross (cross to circle) and respond via button press with the index finger of the right hand. Within each run, 20 stimuli (10 possible and 10 impossible) were presented and repeated 3 times. Three target trials were added for a total of 63 trials per run. The two sessions were identical therefore each of the 120 videos was repeated 6 times (3 repetitions x 2 sessions) across the 12 runs. Additionally, three blank trials were added in each run lengthening the baseline period.

Across sessions, we collected 2 to 3 runs of localizer depending on available scanning time. Each localizer run contained 10 videos per category presented following a block design. Each block lasted 25 seconds (10 videos x 1 sec + 1.5 sec intertrial interval) and was followed by a jittered fixation period of 11 seconds on average. Each category block was repeated 3 times per run. During the localizer participants performed the same task as in the main experiment.

Preprocessing for the functional images was performed using BrainVoyager software (v22.2, Brain Innovation B.V., Maastricht, the Netherlands), Matlab (v2021b) and ANTs (Avants, Tustison, & Song, 2009). To lower thermal noise, we performed NOise reduction with DIstribution Corrected (NORDIC) using both magnitude and phase images (Moeller et al., 2021). EPI Distortion was corrected using the Correction based on Opposite Phase Encoding (COPE) plugin in BrainVoyager, where the amount of distortion is estimated based on volumes acquired with opposite phase-encoding (PE) with respect to the PE direction of the main experiment volumes (Fritz et al., 2014), after which subsequent corrections is applied to the functional volumes. Other preprocessing steps included scan slice time correction using cubic spline, 3D motion correction using trilinear/sinc interpolation and high-pass filtering (GLM Fourier) cut off 3 cycles per run. During the 3D motion correction process, all runs were aligned to the first volume of the first run using the scanner’s intersession auto-align function, ensuring consistent spatial alignment across sessions. Anatomical images were resampled at 0.4mm isotropic resolution using sinc interpolation. To ensure a correct functional-anatomical and functional-functional alignment, the first volume of the first run was coregistered to the anatomical data in native space using boundary based registration (Greve & Fischl, 2009). Functional images were exported in nifti format for further processing in ANTs. To reduce non-linear intersession distortions, functional images were corrected using the antsRegistration command in ANTs using as target image the first volume of the first run and as moving image the first volume of all the other runs. Volume Time Courses (VTCs) were created for each run in the normalized space (sinc interpolation). Prior to the encoding analysis (and following an initial general linear model [GLM] analysis aimed at identifying regions of interest based on the response to the localizer blocks), we performed an additional denoising step of the functional time series by regressing out the stimulus onset (convolved with a canonical hemodynamic response function [HRF]) and the motion parameters. This step was crucial for minimizing the influence of external confounds, such as the timing of stimulus presentation and participant head motion, on the neural data. By removing these factors, we ensured that the model’s training focused exclusively on learning patterns directly associated with the features of the encoding models. However, this approach, while effective in isolating feature-driven neural responses, can lead to smaller accuracies as it also removes some of the variance explained by the stimulation paradigm itself. Despite this trade-off, this method provides a cleaner and more specific evaluation of the encoding models’ ability to capture the relevant neural patterns.

Segmentation of white matter (WM) and gray matter (GM) boundaries as well as cortical layers estimation was performed using a custom pipeline. First, the UNI image and T1 image obtained from MP2RAGE were exported to nifti. We performed gaussian noise reduction using the DenoiseImage command in ANTs (Manjón, Coupé, Martí-Bonmatí, Collins, & Robles, 2010), and bias field correction in SPM12 as described on layer fMRI blog (https://layerfmri.com/2017/12/21/bias-field-correction/). After preprocessing of anatomical images, cortical reconstruction and volumetric segmentation was performed using Basic SAMSEG (cross-sectional processing) command of the Freesurfer image analysis suite (http://surfer.nmr.mgh.harvard.edu/), using the UNI images as T1w contrast and the T1 map of the MP2RAGE (which has flipped intensities between white and gray matter, resembling a T2w image) as T2w contrast. Lastly, cortical thickness and layers extraction were performed using surf_laynii.sh script (https://github.com/srikash/surf_laynii/blob/main/surf_laynii) which enables layering in LAYNII (Huber et al., 2021) using the Freesurfer segmentations output. Three layers were then calculated in LAYNII using the equi-volume approach. All analyses were performed in the individual subject space, but for visualization purposes we projected single-subject statistical or encoding maps onto a group cortex-based aligned surface and then averaged the results across subjects (Goebel, Esposito, & Formisano, 2006).

### Voxel selection for encoding analysis

The functional time series of the localizer runs collected in each participant were analysed using a fixed-effect GLM with 5 predictors (4 conditions in the localizer: Body Objects and their scrambled version and 1 modelling the catch trials). Motion parameters were included in the design matrix as nuisance regressors. The estimated regressor coefficients representing the response to the localizer blocks were used for voxel selection. A voxel was selected for the encoding analysis if significantly active (q(FDR)<0.05) in response to the Body and Objects categories. Note that this selection is unbiased to the response to the stimuli presented in the experimental section of each run.

### Functional ROI definition

Using the functional localizer we also defined body selective regions at the single subject level. Specifically, the EBA was defined using the contrast [Body + Body Scrambled] > [Objects + Objects Scrambled] (Ross, de Gelder, Crabbe, & Grosbras, 2020) with a statistical threshold of q(FDR) < 0.05. All subsequent ROI-level analyses were conducted by identifying the intersection between the voxels assigned to the EBA and those selected for the encoding analysis.

### Encoding models

In order to understand what determines the response to body images we tested several hypotheses, represented by different computational models, using fMRI encoding (Allen et al., 2018; Kay et al., 2008; Naselaris et al., 2011; Santoro et al., 2014). We compared the performance (accuracy in predicting left out data) of three encoding models.

The first model represented body stimuli using the position of joints in three dimensions (kp3d) using 71 keypoints (main skeleton joints like hips, knees, shoulders, elbows, hands and facial features like eyeballs, neck and jaw) extracted from the MoVi dataset. This model represents the stimuli as a collection of points in space forming a human skeleton. To focus on joints that significantly influence perception while minimizing variability from less relevant keypoints, we excluded constant (or almost constant) keypoints ending up with a subset that included 56 keypoints (shoulders, elbows, wrists, hips, knees, and ankles, hands, fingers and facial features from both sides of the body).

The second model quantifies the similarity distances (simdist) between morphed movements (impossible) and normal movements (possible) by analyzing motion capture data extracted during stimulus creation. For each video, both the modified and original motion data were loaded. Initially, all 71 joints defined in the MoVi skeleton were considered. However, to focus on joints with meaningful movement and reduce variability from less relevant joints (such as fingers and toes), joints without rotation data (i.e., joints with empty rotation indices) were excluded, reducing the original set to 56 keypoints (the same as in the previous paragraph). For each selected joint at each time frame, we converted the original Euler angles representing the joint rotation to axis-angle representation. This process yielded a set of three-dimensional vectors in Euclidian space representing the rotation of each joint over time. To measure the similarity between test movements (both modified and original) and the manifold of normal (original) movements, a Gaussian kernel-based approach was employed. This method quantifies the proximity of motion data in the high-dimensional joint angle space, allowing for a robust assessment of movement similarity (see supplementary material). Keypoints for which the computed similarity distances to the normative manifold were not finite (e.g., containing NaN or Inf values) were identified and excluded to maintain data quality, reducing the original 56 keypoints to 29. Similarity distances for all joints were then concatenated to form feature vectors representing each movement’s similarity across all considered joints. This model encoded biomechanical differences because it evaluates the kinematic properties of human joint movements by measuring their distances to a manifold of normal actions, thereby allowing for the differentiation between biomechanically plausible (possible) and implausible (impossible) movements, with the latter exhibiting higher distances due to their deviation from typical human motion patterns. (for the mathematical formulation see supplementary materials). The third model encodes categorical differences between possible and impossible stimuli by incorporating two features that explicitly indicate the (im)possibility of each stimulus. Unlike the other models, this approach does not account for variations within each category, focusing instead on the binary classification of stimuli as either possible or impossible. This model is considered more abstract (or higher-order) compared to the kp3d and simdist models, as it goes beyond image computable approaches (like keypoints) and instead recapitulates a conceptual distinctions.

### Banded ridge regression and model estimates

In the context of fMRI, the linearized encoding framework typically uses L2-regularized (ridge) regression to extract information from brain activity (Hoerl & Kennard, 1970). This method is effective for improving the performance of models with nearly collinear features and helps minimize overfitting. When dealing with multiple encoding models, ridge regression can either estimate parameters for a combined feature space or for each model separately. However, using a single regularization parameter for all models may not be optimal due to varying feature space requirements. To address this, banded ridge regression optimizes separate regularization parameters for each feature space, enhancing model performance by reducing spurious correlations and ignoring non-predictive features.(Dupré la Tour et al., 2022; Nunez-Elizalde et al., 2019). In the present work we used banded ridge regression to fit the three encoding models, combined in a joint encoding model, and performed a decomposition of the variance explained by each of the models following established procedures (Dupré la Tour et al., 2022; Marrazzo et al., 2023).

Model training and testing were performed in cross-validation (3-folds: training on 8 runs [80 stimuli repeated 6 times] and testing on 4 runs [40 repeated 6 times]). For each fold, the training data were additionally split in training set and validation set using split-half crossvalidation. Within the (split-half) training set a combination of random search and gradient descent (Dupré la Tour et al., 2022) was used to optimize the model fit to the data (regularization strength and model parameters). Ultimately, the best model over the two (split-half) validation folds was selected to be tested on the independent test data (4 runs). Within each fold, the models’ representations of the training stimuli were normalized (each feature was standardized to zero mean and unit variance withing the training set). The feature matrices representing the stimuli were then combined with the information of the stimuli onset during the experimental runs. This resulted in an experimental design matrix (nrTRs x NrFeatures) in which each stimulus was described by its representation by each of the models. To account for the hemodynamic response, we delayed each feature of the experimental design matrix (5 delays spanning 11.5 seconds). The same procedure was applied to the test data, with the only difference that when standardizing the model matrices, the mean and standard deviation obtained from the training data were used.

We used banded ridge regression to determine the relationship between the features of the encoding models (stimulus representations) and the fMRI response at each voxel. The encoding was limited to voxels that significantly responded to the localizer stimuli (p(FDR)<0.05) in each individual volunteer’s data. For each cross-validation, we assessed the accuracy of the model in predicting fMRI time series by computing the correlation between the predicted fMRI response to novel stimuli (4 runs, 40 stimuli) and the actual responses. The accuracies obtained across the three folds were Z-transformed and then averaged. To obtain the contribution of each of the models to the overall accuracy we computed the partial correlation between the measured time series and the prediction obtained when considering each of the models individually (Dupré la Tour et al., 2022). Statistical significance was assessed at the group level via permutation test (subject wise sign-flipping, 2^N=2048 times with N=11), and correction for multiple comparison was performed using FDR (q<0.05). Additionally, for each subject, we obtained the average coefficient of determination within EBA (defined in the localizer) and we tested significance with a t-test against zero.

## Results

### Consistent behavioral categorization of possible and impossible stimuli

The analysis of the questionnaire responses showed that all stimuli were accurately categorized. In the "possible" condition, each stimulus received the highest rating, confirming correct classification. Results for the "impossible" videos showed more variability while consistently scoring below 4 on the 1-7 Likert scale. Notably, 95% (57 out of 60) of these stimuli had a median rating between 1 and 2, with the remaining three videos rated between 2 and 3 (see supplementary material for more information).

### Localizer stimuli reveal activation in ventral visual cortex and EBA for voxel selection

In each subject, voxels that significantly responded to the localizer conditions (Body + Objects) with a false discovery rate (FDR) of less than 0.05 were selected for the encoding analysis. While selection took place at the individual level, in Figure 2 we report group-level maps obtained by averaging individual thresholded (q(FDR)<0.05) single-subject maps. All group maps are projected on group aligned (cortex based aligned -CBA) surface. The localizer conditions consistently activated regions in the occipitotemporal cortex, specifically in the superior, middle, and inferior occipital gyri (SOG/MOG/IOG), fusiform gyrus (FG), lingual gyrus (LG) middle temporal gyrus (MTG), inferior temporal sulcus (ITS), lateral occipital sulcus (LOS), and superior temporal sulcus (STS). These clusters overlap with areas identified in our previous study (Marrazzo et al., 2023). By subtracting the responses to object stimuli from the responses to body stimuli, we defined the extrastriate body area (EBA) in each individual and computed probabilistic maps of the overlap of EBA across individuals in cortex based aligned space. The EBA spanned the MOG, MTG, and ITS (Fig. 2) with the probabilistic maps showing an overlap between 20 (white in the Fig.2) and 100% (Green) of subjects.

**Figure 2.**
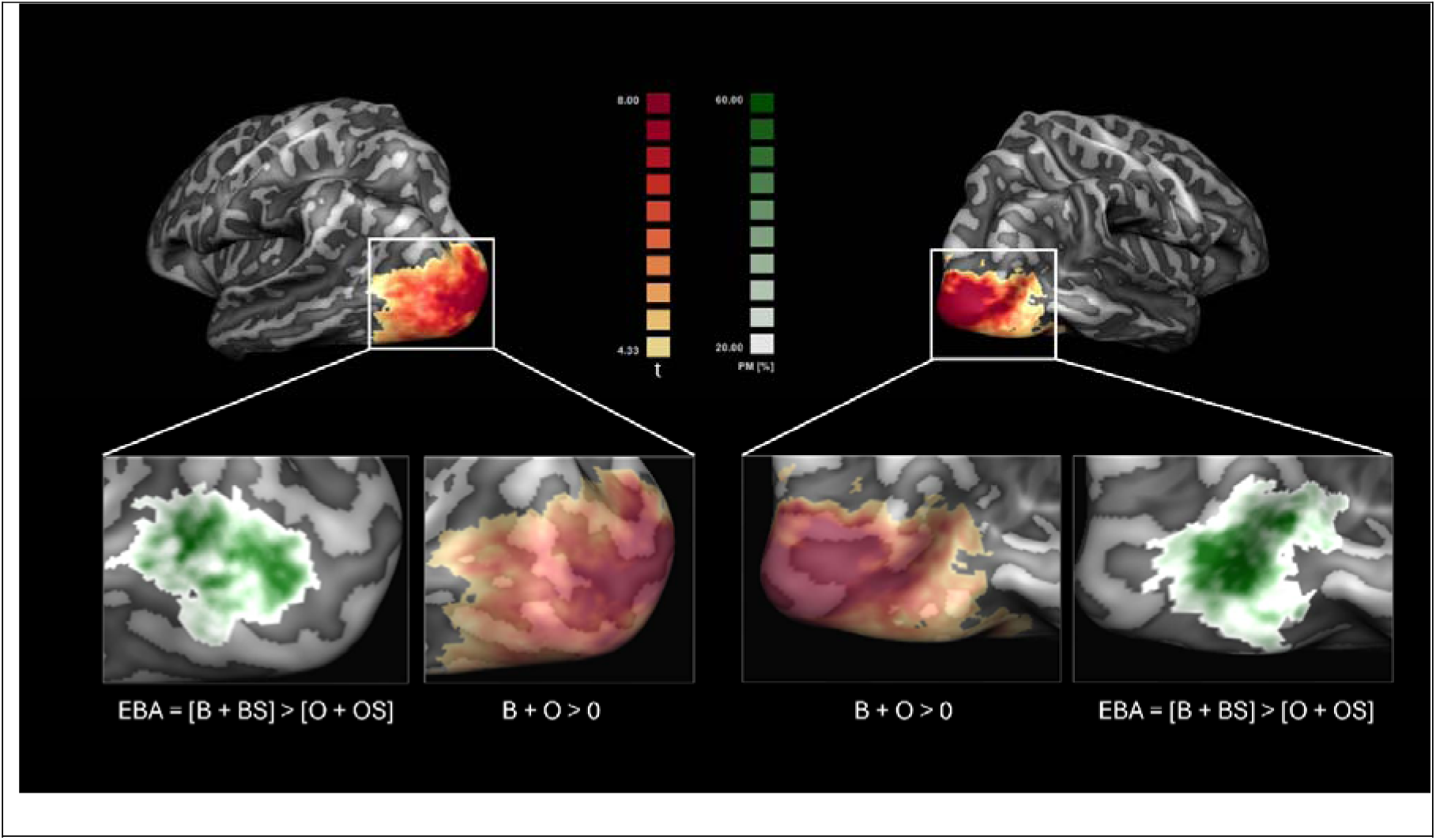
Voxels selection and EBA definition. Voxels that were significantly (q(FDR)<0.05) responding to localizer stimuli [Body + Objects]>0 were selected for the encoding analysis. Although the analysis was performed at single-subject level, for visualization purposes we show the average t-map (in red-yellow) obtained by averaging the thresholded single-subjects maps projected on a group cortex-based aligned mesh. EBA was defined within the localizer via the contrast [Body + Body Scramble] > [Objects + Objects Scramble]. Shown in white-green is a probabilistic map indicating the overlap between individually defined EBAs (q(FDR)<0.05).

### The joint encoding model significantly predicts responses to novel stimuli in ventral visual cortex

The main effect of the responses in the localizer (objects + bodies) was used to select voxels for the encoding in the individual subjects’ data. In these voxels, the response elicited by body stimuli in the main experiment, independent of the localizer, was modelled using banded ridge regression. The group performance of the joint encoding model (kp3d, categorical, simdist) is shown in Figure 3a. Statistical significance at the group level was assessed via a permutation test, with correction for multiple comparisons using FDR (q<0.05). The joint encoding model significantly predicted responses to novel stimuli throughout the ventral visual cortex (SOG, MOG, IOG, ITG, MTG, FG, LOS)

**Figure 3.**
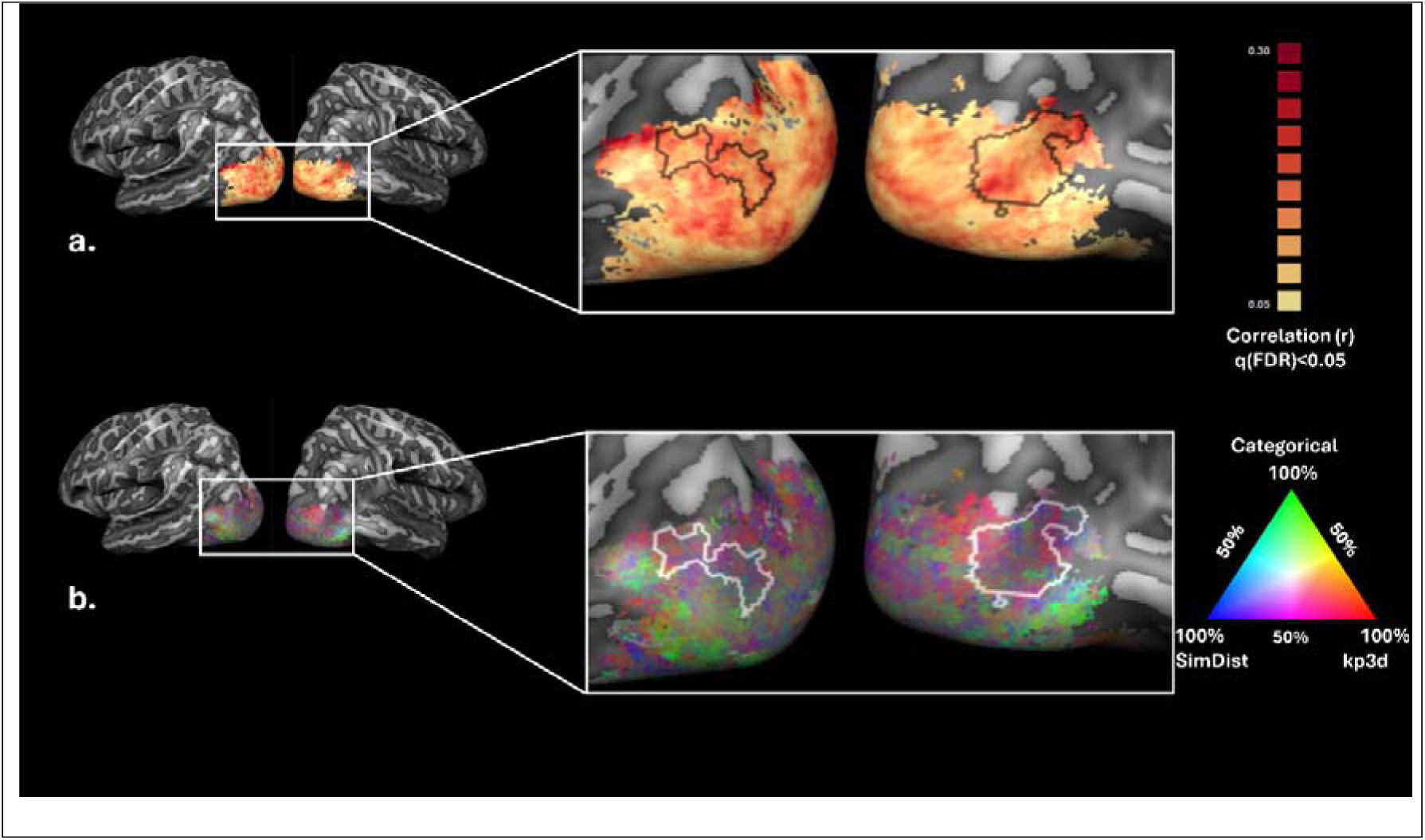
Group-level encoding results. **(a)** Group Prediction accuracy for the joint model (kp3d, categorical, simdist). Statistical significance was assessed via permutation test (subject wise sign-flipping, 2^N=2048 times with N=11), and correction for multiple comparison was performed using FDR (q<0.05). (**b**) RGB map in which each vertex is colour coded according to the relative contribution of each model to the accuracy of the joint model (red = 100% kp3d; blue = 100% simdist; green = 100% categorical) as shown in (a). For clarity, we overlay the outline of EBA as defined in the probabilistic map depicted in Figure 2 by selecting vertices shared by at least 40% of the subjects.

Spatial differences in model contributions to fMRI responses were assessed using an RGB map (Figure 3b), where each vertex is color-coded to show the relative contribution of each model to the joint encoding model’s accuracy. The kp3d model (red) and simdist model (blue) showed varying contributions across regions, with the categorical model (green) also playing a role.

In early visual cortical areas, the response to both possible and impossible bodies was best captured by a combination of the kp3d and simdist models, as indicated by magenta and purple hues. The categorical model (green) contributed more to the voxels’ response in ventral occipital regions, either on its own or in combination with the one of other models (reflected by light-blue or orange colors).

### EBA encodes postural, biomechanical and categorical information

Within the EBA, the joint encoding model accounted for approximately 10% of the variance of the BOLD signal (Fig. 4, top panel). When considering the model fit across cortical layers, we did not observe significant differences in joint model fit across layers, despite a trend for the model fit to increase from inner to superficial layers (Fig. 4 bottom left panels). Although not statistically significant, the percentage of R² explained by each model showed a trend, with the kp3d model accounting for a larger portion of the variance (approximately 40%) (permutation test subject wise sign-flipping, 2^N=2048 times with N=11 on the differences of variance explained: kp3d-simdist, p=0.0698; kp3d-categorical, p=0.083) in the left hemisphere (Fig. 4, bottom right panels). " Moreover, a layer-specific analysis within EBA revealed that the joint model’s performance increased from inner to superficial layers in the right hemisphere (inner-middle: t(10) = -3.546, p = 0.005; inner-superficial: t(10) = -2.325, p = 0.042),

**Figure 4.**
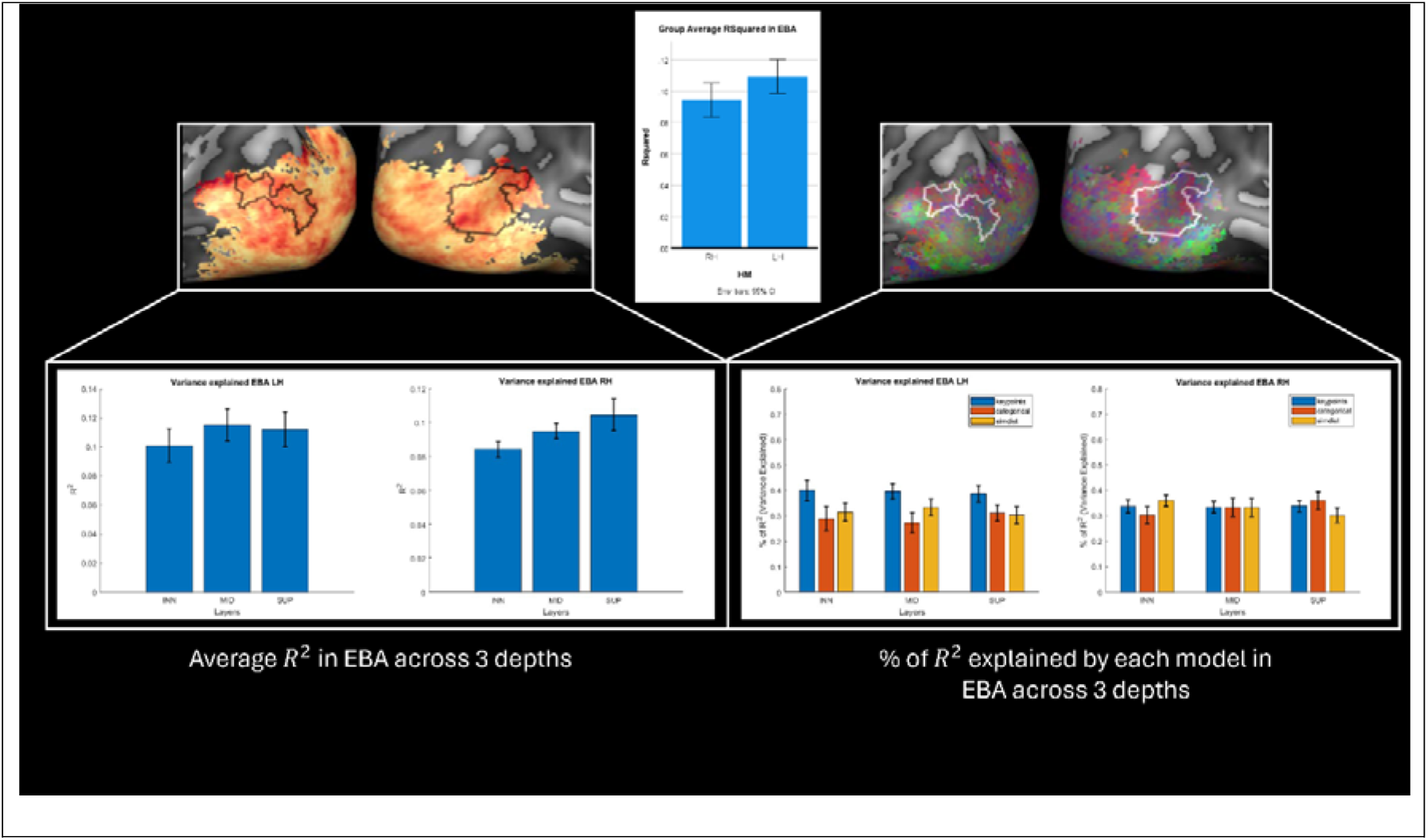
Joint model performance and variance partitioning in EBA across cortical depths. Variance partitioning in the extrastriate body area (EBA) across 11 subjects, comparing left (LH) and right hemispheres (RH) across three cortical layers (Left to right ➔ inner, middle, superficial). The top panel shows the group average R² values in the EBA, indicating overall joint model performance across hemispheres. The bottom left panels display the variance explained (R²) in the LH and RH EBA across layers. The bottom right panel illustrates the percentage of R² explained by each model across layers. To check for differences in variance explained between models, we ran an ANOVA which showed a significant main effect of models (F(2,180) = 4.408, p=0.014) and a significant interaction between hemispheres and models (F(2,180) = 3.572, p=0.030) were found, indicating that models performance varies between hemispheres and that the effectiveness of each model differs across layers. Error bars represent standard errors.

An ANOVA testing for the difference in model performance between hemispheres and layers revealed a significant main effect of models (F=4.408, p=0.014) and a significant interaction between hemispheres and models (F=3.572, p=0.030).

## Discussion

The present study investigated how dynamic body stimuli, specifically biomechanically possible and impossible movements, are encoded in occipitotemporal cortex. Specifically, we compared the predictive performance of encoding models based on 3D keypoints, similarity distances, and categorical differences (kp3d, simdist, categorical). At the group level, we observed that a combination of the three models significantly predicted fMRI BOLD responses in the ventral visual cortex after applying permutation testing and correcting for multiple comparisons. The variance partitioning across the different models of body posture in EBA across cortical layers revealed hemispheric differences between models. In the left hemisphere, the kp3d model appeared to explain a larger portion of the variance (approximately 40%) compared to the simdist and categorical models (both around 30%), with this pattern observed consistently across cortical depths (see Fig. 4). In the right hemisphere, the kp3d model accounted for approximately one-third of the variance, while the simdist and categorical models showed differing trends across cortical layers. Specifically, the simdist model tended to explain more variance in the inner layers, whereas the categorical model appeared to account for more variance in the superficial layers. Although these differences were not statistically significant.

### Low-level and high-level features in the occipitotemporal cortex

Our findings reveal that a combination of low and high-level features contribute to the dynamic perception of body movement in occipitotemporal cortex. In early visual cortical areas, the kp3d and simdist models alone, or in combination, best predicted neural responses (red, blue, magenta-purple color patches in Fig. 3b) indicating that postural and biomechanical features play a significant role in these regions. These results align with the notion that early visual areas process low-level features such as orientation, spatial frequency, and basic shape attributes (Carandini et al., 2005; Kay et al., 2008; Naselaris et al., 2011; Nishimoto & Gallant, 2011; Nishimoto et al., 2011). As processing advances to higher visual areas, the categorical model becomes increasingly dominant. This shift aligns with previous literature showing that higher-order areas integrate lower-level features into more abstract representations, reflecting a progression toward semantic processing (Grill-Spector & Weiner, 2014; Haxby et al., 2001; Huth et al., 2012; Kriegeskorte, Mur, & Bandettini, 2008).

### Encoding of body stimuli in EBA

Within the EBA, our analysis revealed that a combination of the three encoding models— kp3d, simdist, and categorical—significantly predicted neural responses, accounting for approximately 10% of the variance of the BOLD signal in EBA (see Fig. 4). This indicates that in EBA these various types of information, including postural and biomechanical features and categorical distinctions are combined.

While all models contributed significantly to the response elicited by dynamic bodies in EBA, this was more prominent for the kp3d and simdist models (purple-magenta patches – in Fig. 3b and 4) in the superior part of EBA, covering middle occipital gyrus (MOG) and superior occipital gyrus (SOG). In contrast, the anterior inferior part of EBA —spanning anterior part of the inferior temporal gyrus (aITG) and anterior lateral occipital sulcus (aLOS)— tended towards categorical encoding (cyan-orange-green patches in Fig. 3b and 4) suggesting an integration of postural information in the keypoint model with more abstract representations. This may involve linking specific body configurations to semantic information such as the type of action being performed or the emotional state conveyed by the body movement (Foster et al., 2021; Foster et al., 2019)

This functional heterogeneity found in EBA aligns with anatomical findings that identify distinct body-selective areas within the occipitotemporal cortex (Weiner & Grill-Spector, 2011). Recent findings by Li et al. (Li, Poyo Solanas, Marrazzo, & de Gelder, 2024) using data-driven methods identified four adjacent body-selective nodes within the occipitotemporal cortex further support this notion. Specifically, the predominance of kp3d and simdist in superior subregions may reflect their role in detailed sensory processing, as they show stronger connectivity with regions involved in processing fine-grained visual details (Li et al., 2024). In contrast, the anterior inferior subregions’ reliance on categorical encoding suggests involvement in higher-order interpretation and integration of body-related information, consistent with their broader connectivity profiles (Li et al., 2024). Our findings thus reinforce the notion that EBA is functionally heterogeneous consistent with the finding of specialized subregions dedicated to different aspects of body and action perception. (Li et al., 2024).

Furthermore, our results are consistent with previous findings showing that EBA is more functionally and structurally connected to dorsal stream regions compared to other body-related areas, such as FBA and the lateral occipital complex (LOC) (Zimmermann, Mars, de Lange, Toni, & Verhagen, 2018). This connectivity supports the EBA’s role in bridging perceptual and motor functions, particularly in specifying goal-directed postural configurations for motor planning. Notably, the study suggests that EBA’s connectivity with parietal regions, such as the superior parietal lobule and postcentral gyrus, may enable it to access somatosensory information, which is essential for planning and executing actions based on body information. This suggestion is consistent with the earlier findings from (Astafiev, Stanley, Shulman, & Corbetta, 2004) reporting that the EBA responds to goal directed movements of the observers’ body parts.

### Layer-specific encoding in EBA

Our layer-specific analysis within EBA revealed that the joint model’s performance increased from inner to superficial layers in the right hemisphere (inner-middle: t(10) = -3.546, p = 0.005; inner-superficial: t(10) = -2.325, p = 0.042), which may hint towards a gradient of sensitivity to postural features in right EBA. Conversely, the joint model performed uniformly across layers in the left hemisphere. The variance partitioning also hinted at a potential hemispheric difference, with the kp3d model accounting for a substantial portion of the variance (approximately 40%) across all cortical depths in the left hemisphere. This trend may suggest a specialization for encoding detailed three-dimensional postural information., which is essential for precise spatial judgments and the accurate interpretation of body movements (Caspari et al., 2014; Kumar, Popivanov, & Vogels, 2019). This left-lateralization aligns with previous findings indicating a dominance of the left hemisphere in processing detailed aspects of body stimuli (Bracci et al., 2010; Downing & Peelen, 2016) and may enhance the ability to recognize and interpret fine-grained body movements, facilitating action recognition and understanding others’ intentions. (Blake & Shiffrar, 2007; de Gelder et al., 2010; Urgesi, Candidi, Ionta, & Aglioti, 2007).

In the right hemisphere, we observed a varying contributions trend of the simdist and categorical models across cortical layers—higher simdist influence in inner layers and greater categorical influence in superficial layers— which may hint at a layer-specific encoding strategy. Although these differences were not statistically significant, they hint at a potentially differentiated role of cortical layers in processing biomechanical and categorical information. This aligns with the notion that deeper cortical layers may handle more input-driven, sensory information, while superficial layers integrate higher-order, contextual, or semantic information (Bastos et al., 2012; Felleman & Van Essen, 1991; Larkum, 2013; Rockland & Pandya, 1979; Spratling, 2017). Additionally, a similar depth-dependent organization has been demonstrated in the ventral temporal cortex, where superficial layers predominantly encoded broader, domain-level distinctions, while deeper layers were more sensitive to specific category-level information (Margalit et al., 2020).

### Role of biomechanical plausibility

The substantial predictive power of the simdist model from early to high-level visual cortex underscores the visual system’s sensitivity to biomechanical constraints from the initial stages of processing. This suggests that, beyond simply recognizing body parts, the brain may be encoding midlevel features (de Gelder & Poyo Solanas, 2021) that reflect the biomechanical characteristics of human bodies. Midlevel features, including biomechanical constraints on posture and movement, could serve as a crucial link between the configurations driven by body joints we previously identified (see (Marrazzo et al., 2023)) and more abstract, higher-level representations of the body.

Furthermore, differentiating between possible and impossible movements likely involves detecting deviations from typical joint configurations and movement patterns, which could suggest the presence of an internal model of human biomechanics (P. Dayan & Berridge, 2014; Giese & Poggio, 2003). Our results extend previous findings by demonstrating that encoding models can effectively capture neural responses to biomechanical plausibility (Costantini et al., 2005). This sensitivity may reflect a mechanism for detecting errors or anomalies in observed body movement, acting as a filtering mechanism for upstream processing of actions, which is critical for social cognition (Candidi, Urgesi, Ionta, & Aglioti, 2008; Kilner, Friston, & Frith, 2007; Li et al., 2024; Schubotz, 2007; Urgesi et al., 2007).

### Limitations and future directions

Our scanning parameters focused primarily on occipitotemporal and frontal regions, excluding areas such as the motor and premotor cortices. These regions are known to play a crucial role in the recognition of both static and dynamic bodily actions (Pobric & de C. Hamilton, 2006; Urgesi et al., 2007), responding to biomechanically possible and impossible stimuli alike (Costantini et al., 2005) and contributing to the distinction between actions that can be performed and those that cannot (Candidi et al., 2008). Incorporating these regions in future studies will help clarify their role in the perception and discrimination of biomechanical plausibility, offering a more comprehensive view of the neural mechanisms underlying action recognition. Additionally, our stimulus creation was limited to manipulations of the elbows and knees to generate impossible movements. Future research might include a broader range of movements and joint manipulations to evaluate the generality of encoding mechanisms across different biomechanical contexts. Also, the relatively small sample size (n=11) is common in laminar fMRI studies, but may limit the generalizability of our findings. Replication with larger samples is needed to confirm the observed effects and strengthen the reliability of these results. Finally, further exploration of hemispheric differences, along with the potential influence of attention and task demands on encoding, would enrich our understanding of the factors shaping these neural processes.

## Conclusions

In summary, this study investigated whether occipitotemporal cortex, particularly the body sensitive area EBA, encodes biomechanically possible and impossible body movements. By comparing three encoding models—3D keypoints, similarity distances, and categorical differences—we found that a combination of these models significantly predicted neural responses in the ventral visual cortex. In the left hemisphere, 3D keypoints explained a larger portion of the variance across cortical layers, while in the right hemisphere we saw an emerging trend for preference for similarity distances in deeper layers, with categorical differences becoming more prominent in superficial layers. The study underscores the brain’s sensitivity to biomechanical plausibility, with the biomechanical (simdist) model explaining a significant portion of the variance from early stages of visual processing. Lastly, these findings highlight the EBA’s functional heterogeneity, with superior regions (middle/superior occipital gyri) focusing on detailed biomechanical features and anterior regions (lateral occipital sulcus and inferior temporal gyrus) integrating more abstract, categorical information.

## Acknowledgements

This work was supported by the European Research Council (ERC) Synergy grant (Grant agreement 856495 Relevance), by the Horizon 2020 Programme H2020-FETPROACT430 2020-2 (Grant agreement 101017884 GuestXR) and Horizon 2020 grant (Grant agreement 101070278 ReSilence).

## Data availability statement

Data and code are being prepared for public availability.

## CRediT authorship contribution statement

**Giuseppe Marrazzo:** Conceptualization, Investigation, Software, Formal analysis, Validation, Visualization, Writing – Original Draft Preparation, Writing -review & editing. **Federico De Martino:** Conceptualization, Investigation, Supervision, Validation, Writing -review & editing. **Albert Mukovskiy:** Software, Validation, Writing -review & editing. **Martin A. Giese:** Conceptualization, review & editing, Funding Acquisition. **Beatrice de Gelder:** Conceptualization, Project Administration, Resources, Supervision, Funding Acquisition, Writing – Original Draft Preparation, Writing -review & editing

