## Supplementary Material for "Neural encoding of biomechanically (im)possible human movements in occipitotemporal cortex" (from another version of this preprint: version 1, 2025-01-07)

#### 1 Material and Methods

##### 1.1 Biomechanical model (simdist)

To analyze the similarity between morphed movements and normal movements, we utilized motion capture data corresponding to the stimulus videos used in our experiment. The dataset comprised both modified movements (morphed movements) and original, unmodified movements. To measure the similarity between test movements (both modified and original) and the manifold of normal (original) movements, a Gaussian kernel-based approach was employed. This method quantifies the proximity of motion data in the high-dimensional joint angle space, allowing for a robust assessment of movement similarity. The axis-angle data from all time frames and all original (normal) movements were concatenated to form a comprehensive dataset representing the normal actions manifold. This manifold encompasses the typical joint configurations observed during natural movements across all subjects and actions in the experiment.

##### 1.2 Estimation of the Gaussian Width Parameter

To measure similarity in the high-dimensional joint angle space, we employed a Gaussian kernel that required estimation of the Gaussian width parameter $\sigma$. This parameter was crucial for appropriately scaling distances within the kernel function.

##### 1.3 Local variance estimation

For each point $x_n$ in the normal actions manifold, we identified its $N = 8$ nearest neighbors based on Euclidean distance in the joint angle space. Due to the high dimensionality of the data (each point represents the axis-angle components of multiple joints), we utilized a k-d tree algorithm (Ranger) (see `https://www3.`

cs.stonybrook.edu/~algorith/implement/ranger/implement.shtml) for efficient nearest neighbor searches. We calculated the average squared distance to the nearest neighbors for each point to estimate the local density:

$$\mathrm{dist}(n) = \frac{1}{N}\sum_{i=1}^{N} \|x_n - x_{\mathrm{n,i}}\|^2 \tag{1}$$

where $x_{\mathrm{n,i}}$ represents the $i$-th nearest neighbor of point $x_n$.

The Gaussian width parameter $\sigma^2$ was calculated by taking the average of these local distances across all points in the manifold:

$$\sigma^2 = \mathrm{mean}(\mathrm{dist}(n)) \tag{2}$$

This estimation ensures that $\sigma$ reflects the typical local variance within the data, allowing the Gaussian kernel to appropriately weigh the contributions of different points based on their proximity.

##### 1.4 Similarity Measure Calculation

To quantify the similarity between each test movement (both modified and original) and the normal actions manifold, we utilized a Gaussian kernel-based similarity measure.

For a test movement with $T$ frames and joint data $\{x_{\mathrm{test}}(t)\}_{t=1}^{T}$, the similarity measure $S$ was computed as:

$$S = \sum_{n=1}^{N_{\mathrm{orig}}} \sum_{t=1}^{T} \exp\left(-\frac{\|x_{\mathrm{test}}(t) - x_n\|^2}{2\sigma^2}\right) \tag{3}$$

where:

- $N_{\mathrm{orig}}$ is the number of points in the normal actions manifold, namely all time points in all normal actions trajectories.
- $x_n$ are the points in the normal actions manifold.
- $\|\cdot\|$ denotes the Euclidean norm in the joint angle space and $\sigma$ as in (2).

To normalize the similarity measure and transform it into a distance metric, we defined the similarity distance as:

$$\mathrm{simdist} = -\log\left(\frac{S}{N_{\mathrm{orig}} \times T}\right) \tag{4}$$

This formulation yields a positive value where lower similarity distances indicate a higher similarity between the test movement and the normal actions manifold.

### 2 Results

#### 2.1 Behavioral analysis

The analysis of the questionnaire responses showed that for the "possible" stimuli, 98% (59 out of 60) were recognized as daily actions, with median ratings between 6 and 7. Conversely, 93% of the "impossible" stimuli (56 out of 60) were not associated with any action, receiving median ratings between 1 and 3. The remaining four stimuli had a median rating of 4. Participants reported focusing on limb movements and parts (arms, legs, hands or feet) in 96.67% of the "impossible" videos (58 out of 60), while 3.33% (2 out of 60) of the videos focused on overall body movement. In contrast, the "possible" videos showed an increased focus on overall body movement (16.67%, 10 out of 60), though the majority still concentrated on limb movements and parts (83.33%, 50 out of 60).
